# Genetic variability in COVID-19-related genes in the Brazilian population

**DOI:** 10.1101/2020.12.04.411736

**Authors:** Rodrigo Secolin, Tânia K. de Araujo, Marina C. Gonsales, Cristiane S. Rocha, Michel Naslavsky, Luiz De Marco, Maria A.C. Bicalho, Vinicius L. Vazquez, Mayana Zatz, Wilson A. Silva, Iscia Lopes-Cendes

**Affiliations:** Department of Translational Medicine, University of Campinas (UNICAMP), and the Brazilian Institute of Neuroscience and Neurotechnology (BRAINN), Campinas, SP, Brazil; Departament of Genetics and Evolutive Biology, Institute of Bioscience, University of São Paulo, (USP) and the Human Genome and Stem Cell Research Center, São Paulo, SP, Brazil; Department of Surgery, Federal University of Minas Gerais (UFMG), Belo Horizonte, MG, Brazil; Department of Clinical Medicine, Federal University of Minas Gerais (UFMG), Belo Horizonte, MG, Brazil; Molecular Oncology Research Center (CPOM), Barretos Cancer Hospital, Barretos, SP, Brazil; Department of Genetics, Ribeirão Preto Medical School, University of São Paulo at Ribeirão Preto (USP), Ribeirão Preto, SP, Brazil

**Author notes:** **Correspondence to:** Iscia Lopes-Cendes, M.D., Ph.D., Professor of Medical Genetics and Genomic Medicine, Department of Translational Medicine, School of Medical Sciences, University of Campinas (UNICAMP), Tessália Vieira de Camargo, 126., Campinas, SP, Brazil 13083-887. These authors contribute equally to this work (Rodrigo Secolin, Tânia K. de Araujo).

**Keywords:** population genomics, admixed population, COVID-19, SARS-Cov-2

## Abstract

SARS-CoV-2 employs the angiotensin-converting enzyme 2 (ACE2) receptor and the transmembrane serine protease (TMPRSS2) to infect human lung cells. Previous studies have suggested that different host genetic backgrounds in *ACE2* and *TMPRSS2* could contribute to differences in the rate of infection or severity of COVID-19. Recent studies also showed that variants in 15 genes related to type I interferon immunity to influenza virus could predispose to life-threatening COVID-19 pneumonia. Additional genes (*SLC6A20, LZTFL1, CCR9, FYCO1, CXCR6, XCR1, IL6, CTSL, ABO*, and *FURIN*) and HLA alleles have also been implicated in response to infection with SARS-CoV-2. Currently, Brazil has recorded the third-highest number of COVID-19 patients worldwide. We aim to investigate the genetic variation present in COVID-19-related genes in the Brazilian population. We analysed 27 candidate genes and HLA alleles in 954 admixed Brazilian exomes. We used the information available in two public databases (http://www.bipmed.org and http://abraom.ib.usp.br/), and additional exomes from individuals born in southeast Brazil, the region with the highest number of COVID-19 patients in the country. Variant allele frequencies were compared with the 1000 Genomes Project phase 3 (1KGP) and the gnomAD databases. We found 395 non-synonymous variants; of these, 325 were also found in the 1000 Genome Project phase 3 (1KGP) and/or gnomAD. Six of these variants were previously reported as putatively influencing the rate of infection or clinical prognosis for COVID-19. The remaining 70 variants were identified exclusively in the Brazilian sample, with a mean allele frequency of 0.0025. *In silico* prediction of the impact in protein function revealed that three of these rare variants were pathogenic. Furthermore, we identified HLA alleles that were previously associated with COVID-19 response at loci DQB1 and DRB1. Our results showed genetic variability common to other populations, but also rare and ultra-rare variants exclusively found in the Brazilian population. These findings could potentially lead to differences in the rate of infection or response to infection by SARS-CoV-2 and should be further investigated in patients with the disease.

## Introduction

COVID-19 disease, caused by the SARS-CoV-2 coronavirus, is currently a worldwide pandemic. To enter human lung cells, SARS-CoV-2 employs the spike protein, which is primed by the host serine protease (TMPRSS2), followed by angiotensin-converting enzyme 2 (ACE2) receptor binding, and proteolysis with activation of membrane fusion within endosomes by cathepsin L (CTSL)^1-4^. The main feature in SARS-CoV-2 infection is pre-activation of the spike protein by FURIN inside the host cell, which leads to increased SARS-CoV-2 spread into lung cells and increased virulence^5^. The rapid SARS-CoV-2 infection leads to an exacerbated immune reaction, and a few studies have shown that increased levels of IL-6 (an essential immune response mediator) are associated with increased inflammatory response, respiratory failure, increased probability of intubation, the presence of clinical complications, and higher mortality in patients with COVID-19^6-8^. Additional studies found the enrichment of rare variants predicted to be loss-of-function in genes related to type I interferon (IFN) immunity to influenza virus among patients with life-threatening COVID-19 pneumonia (*TLR3, TICAM1, TRIF, UNC93B1, TRAF3, TBK1, IRF3, NEMO, IKBKG, IFNAR1, IFNAR2, STAT1, STAT2, IRF7*, and *IRF9*)^9^.

Specific variants in the *ACE2* and *TMPRSS2* genes have been reported among diverse populations worldwide, suggesting that different host genetic backgrounds could contribute to differences in COVID-19 infection and severity^2,10^. Ellinghaus et al.^11^ performed a genome-wide association study (GWAS) including Italian and Spanish patients with confirmed COVID-19 and controls and identified six candidate genes associated with COVID-19 response on chromosome (chr) 3p21.31 (*SLC6A20, LZTFL1, FYCO1, CXCR6, XCR1, CCR9*), and one on chr 9q34.2, the locus for genes encoding for the *ABO* blood group antigens. A subsequent, more extensive study replicated the association between the locus on chr 3p21.31 and COVID-19. It revealed a COVID-19 risk core haplotype ranging from 45,859,651bp to 45,909,024bp, which was inherited from Neanderthals and is currently carried by approximately 50% of people in South Asia and about 16% of people in Europe^12^. Interestingly, no evidence of association was found for the previously identified candidate genes that are potentially involved in the response to infection by SARS-CoV-2: *ACE2, TMPRSS2, FURIN*, and *IL6*.

Furthermore, one significant factor modulating resistance or susceptibility to viral infections is the human leukocyte antigens (HLAs). HLA polymorphism results from a set of amino-acid substitutions in the peptide-binding groove of the HLA molecules that produce variability in the peptide epitope binding-site and presentation to T cells, which may protect against epidemic infection^13^. Thus, genetic variability in the HLA alleles could influence the immune response in patients with COVID-19, modulating disease severity. Indeed, *in silico* analysis found that HLA-B*46:01 had the fewest predicted binding sites for SARS-CoV-2 peptides, and HLA-B*15:03 showed the greatest capacity to present highly conserved shared SARS-CoV-2 peptides to immune cells^14^.

Brazil has reported the third-highest number of COVID-19 infections worldwide (updated on September 28^th^ 2020; https://covid19.who.int/; https://coronavirus.jhu.edu/map.html), and the highest number of cases is concentrated in the south-eastern region of the country (updated on September 28^th^ 2020; https://covid.saude.gov.br/). Brazilian individuals feature an admixed genome, encompassing European, sub-Saharan African, and Native Americans as the three main ancestry populations^15-17^, and the distribution of ancestry components varies remarkably throughout the genome^18^. Furthermore, it has been demonstrated that a significant proportion of genetic variability is still undiscovered in admixed Brazilians^19^ and that genetic variability may lead to differential response to infection^20^. Therefore, we aim to investigate the genetic variation present in COVID-19 related genes in the Brazilian population.

## Results

### Exome analysis

We found 7,172 variants among the candidate genes analysed (Supplementary Table 1). Of these, 395 variants putatively impact protein function, including 354 non-synonymous variants, seven frameshift substitutions, three in-frame deletions, one in-frame insertion, 12 stop gains, two start losses, and 16 splice site variants (Supplementary Table 1). Three hundred and twenty-five variants were also present in the gnomAD and/or 1000 Genome Project phase 3 (1KGP) databases, including 56 common variants, with an alternative allele frequency (AAF) ≥0.01 and 269 rare variants (AAF<0.01) (Supplementary Data 1). Although AAF from the admixed Brazilian sample follows the distribution from NFE/EUR, AFR, and AMR in gnomAD and 1KGP databases (Fig. 1), we found differences in the AAF of these common and rare variants in the admixed Brazilian sample compared to gnomAD^21^ and/or 1KGP^22^ databases shown in Fig. 1 and Supplementary Data 1. Interestingly, we also observed some variability in the AAF among samples from different Brazilian towns and the two public databases of genomic information on the Brazilian population, BIPMed and ABraOM (Fig. 2).

**Figure 1.**
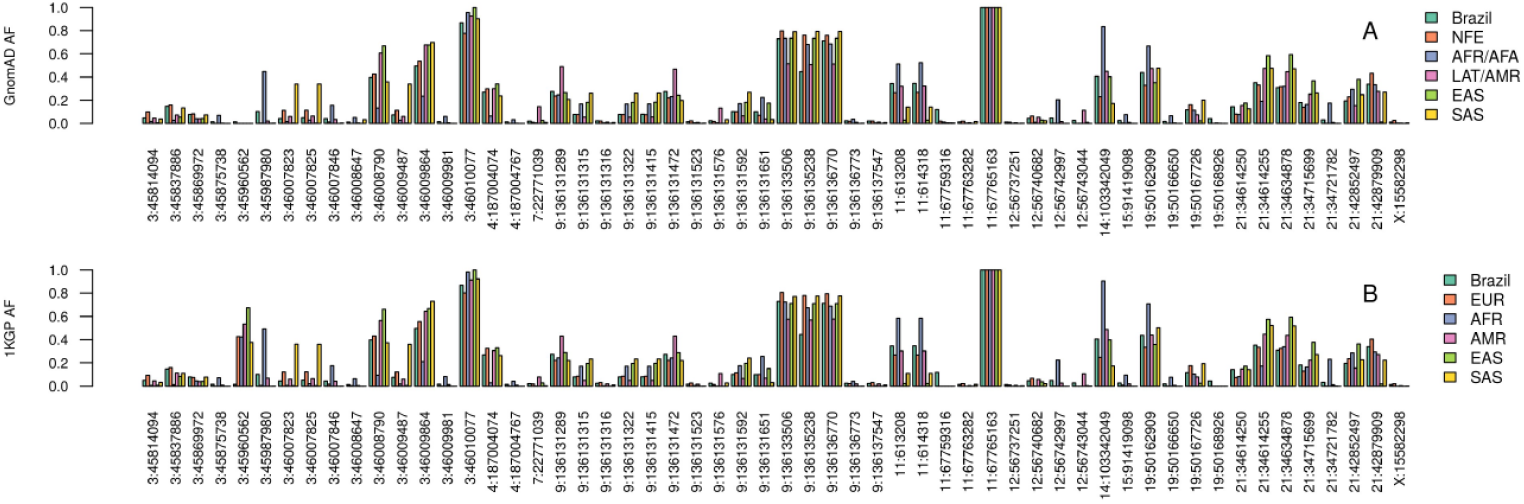
Distribution of alternative allele frequency of common variants (AAF≥0.01) from samples of admixed Brazilians and worldwide public datasets. The x-axis shows the 56 variants found in common between the Brazil sample and the gnomAD and 1KGP dataset. (A) Comparison between Brazilians and gnomAD, and (B) including non-Finland Europeans (NFE), sub-Saharan Africans/African Americans (AFR) Venn diagrams that show the overlap between samples (A) and variants (B) in the WES and the SNP array datasets from BIPMed reference samples.

**Figure 2.**
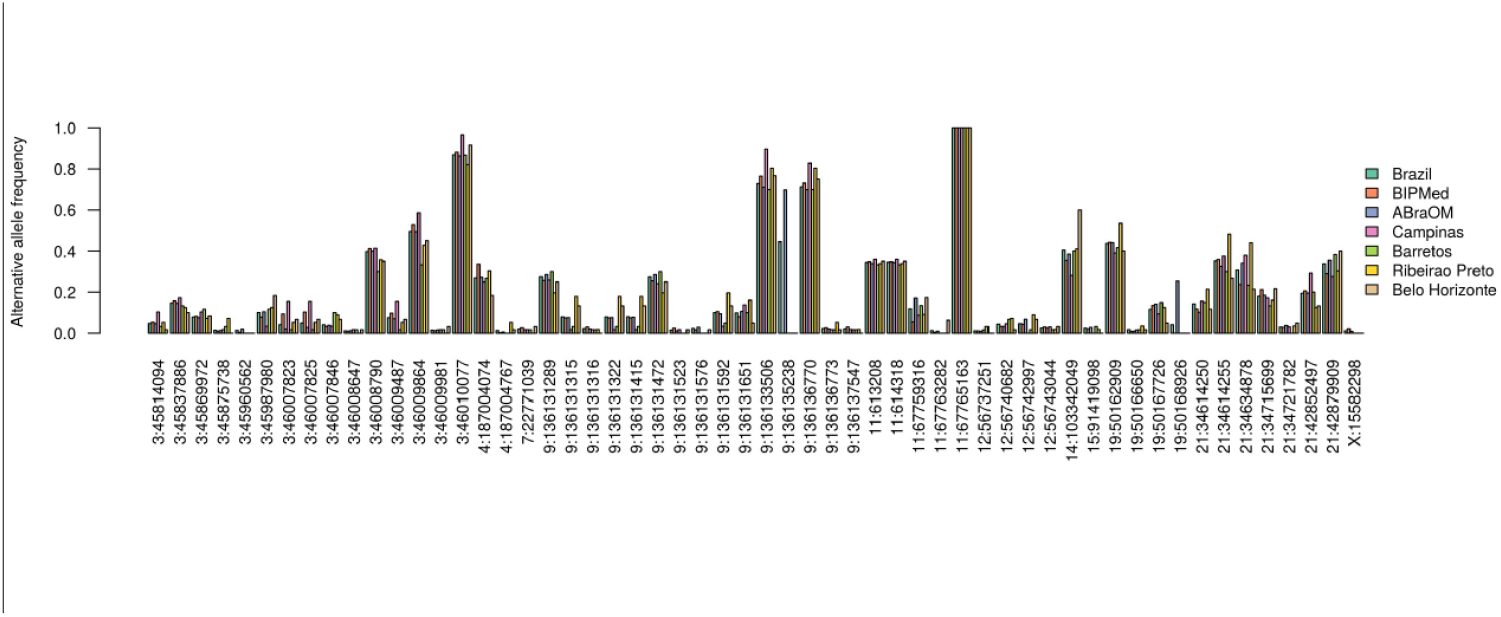
Distribution of alternative allele frequency of common variants (AAF>0.01) separated by Brazilian cities. The x-axis shows the 56 variants found in common with gnomAD and 1KGP. This barplot also includes the two public Brazilian datasets (BIPMed and ABraOM) and the frequency of all samples combined (Brazil).

More importantly, there were 70 variants which were exclusive to the Brazilian sample, including 11 variants in genes related to type I INF immunity to influenza virus^9^, six in candidate genes for COVID response identified by GWAS^11^, and five related to SARS-CoV-2 entry in lung cells and virus replication^2,10^. These are rare or ultra-rare variants, presenting a mean AF of 0.0025 (Supplementary Data 1). Among these, we found one in the dataset from Belo Horizonte and two in the ABraOM database for *ACE2* p.Arg219Cys; one in the dataset from Barretos and two in the ABraOM database for *ACE2* p.Leu731Phe; and the *TMPRSS2* p.Val160Met variant was present in samples from all the different Brazilian towns and the two public databases (BIPMed and ABraOM), with an AAF ranging from 0.1333 in Belo Horizonte to 0.2931 in Campinas. Among the reported variants in genes influencing type I INF immunity to influenza virus^9^, we found three variants in the ABraOM database (one *TLR3* p.Pro554Ser, one *IFR3* p.Asn146Lys and one *IRF7* p.Pro246Ser) (Supplementary Data1 and 2).

In addition, we identified five variants (rs35044562, rs34326463, rs35508621, rs67959919, and rs35624553) which were previously described in the COVID-19 risk core haplotype and inherited from Neanderthals^12^. These were only present in samples from Ribeirão Preto and the BIPMed dataset (rs34326463), Campinas (rs35044562, and rs35508621), and the ABraOM dataset (rs35044562, rs35508621, rs67959919, and rs35624553) (Table 1).

**Table 1.** Alternative allele frequency (AAF) of variants encompassing the COVID-19 risk core haplotype and the alleles present in Neanderthal samples.

| dbSNP | Chr | Pos | ALT | Brazil AF | Risk allele* | Campinas | Barretos | Ribeirão Preto | Belo Horizonte | BIPMed | ABraOM |
| --- | --- | --- | --- | --- | --- | --- | --- | --- | --- | --- | --- |
| rs35044562 | 3 | 45909024 | G | 0.0011 | G | 0.0862 | 0.0000 | 0.0000 | 0.0000 | 0.0279 | 0.0311 |
| rs34326463 | 3 | 45899651 | G | 0.0011 | G | 0.0000 | 0.0000 | 0.0357 | 0.0000 | 0.0000 | 0.0000 |
| rs35508621 | 3 | 45880481 | C | 0.0011 | C | 0.0345 | 0.0000 | 0.0000 | 0.0000 | 0.0000 | 0.0039 |
| rs67959919 | 3 | 45871908 | A | 0.0011 | A | 0.0000 | 0.0000 | 0.0000 | 0.0000 | 0.0000 | 0.0039 |
| rs35624553 | 3 | 45867440 | G | 0.0262 | G | 0.0000 | 0.0000 | 0.0000 | 0.0000 | 0.0000 | 0.0039 |

\*data extracted from Zeberg and Pääbo, 2020 (doi:10.1038/s41586-020-2818-3)

### In silico predictions

We identified seven variants that were predicted to affect protein function for the 12 algorithms used: p.Phe249Ser, p.Gly164Val, and p.Leu25Pro in *the SLC6A20* gene; p.Leu96Arg in *LZTFL1*; p.Tyr287Ser in *XCR1*; and p.Gly146Ser and p.Asn414Ser in the *FURIN* gene (Table 2). Furthermore, the variant p.Gly146Ser in the *FURIN* gene was predicted to destabilise the protein (ΔΔG: −1.576 kcal/mol). We observed that the p.Phe249Ser variant is present in samples from Barretos, the BIPMed dataset, gnomAD, and 1KGP (NFE/EUR, AFR, AMR, and SAS populations), whereas the p.Gly164Val variant is present in the ABraOM dataset, gnomAD, and 1KGP (NFE/EUR populations), and the p.Gly146Ser variant is present in the ABraOM dataset, gnomAD, and 1KGP (NFE/EUR, AFR, AMR, EAS, and SAS populations). Notably, four of the variants predicted to be deleterious are found exclusively in admixed Brazilian individuals (p.Leu25Pro in Barretos; p.Leu96Arg in the ABraOM dataset; p.Tyr287Ser in Belo Horizonte; and p.Asn414Ser in the BIPMed dataset).

**Table 2.**
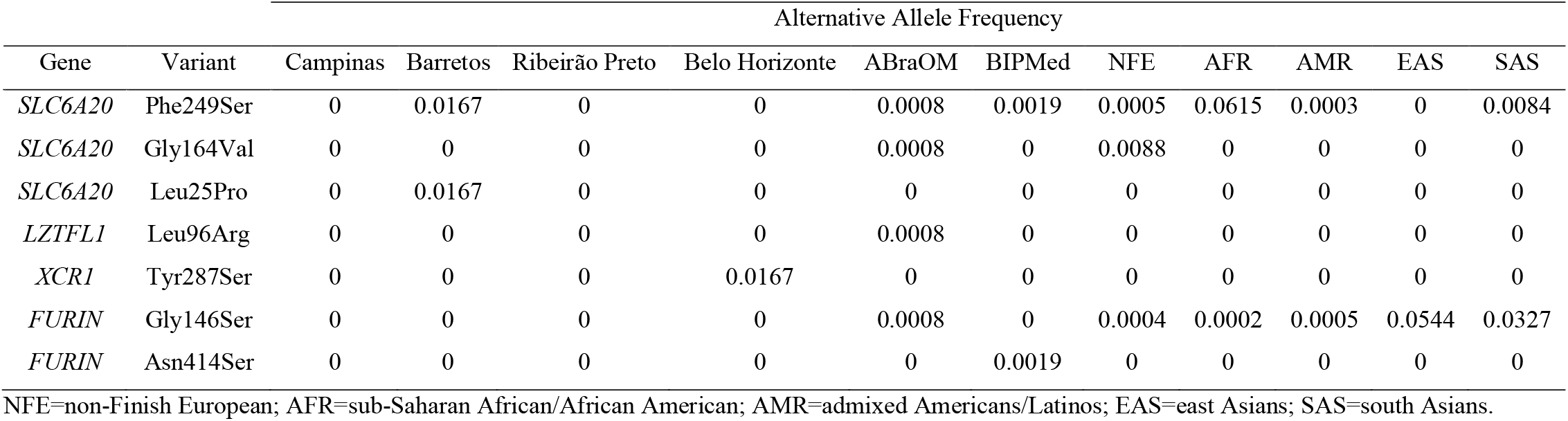
Alternative allele frequency of deleterious variants according to 12 different prediction algorithms

We did not find any predicted deleterious variants in *ACE2* and *TMPRSS2* based on our 12 algorithm criteria. However, Hou et al.^2^ applied only Polyphen2 and CADD scores to variants in *ACE2* and *TMPRSS2* (Polyphen2 >0.96 and CADD >20 as the cut-off). Therefore, only variants defined as ‘probably damaging’ by Polyphen2 (http://genetics.bwh.harvard.edu/pph2/dokuwiki/overview) and CADD (>20) were included. We found 79 variants predicted to affect protein function, including the p.Val160Met variant in *TMPRSS2* reported by Hou et al.^2^, and the p.Pro554Ser variant in *TLR3* previously reported by Zhang et al.^9^ (Supplementary Data 2).

### HLA analysis

Overall, we identified 331 different HLA alleles in the admixed Brazilian samples. Of these, three HLA alleles have been previously associated with COVID-19 response^14,23^. We compared the frequency of these HLA alleles in admixed Brazilians and in populations that occupy the top 10 positions with most cases of COVID-19 and the five populations less affected by the disease, including the United States, India, Russia, Colombia, Peru, Mexico, Spain, Argentina, South Africa, Japan, Australia, South Korea, Vietnam, and Taiwan. The frequency of these alleles is described in Supplementary Data 3. We noticed that the HLA-B*46:01, HLA-B*27:07, HLA-B*15:27, and HLA-C*07:29 alleles were absent in the Brazilian samples. The HLA-C*07:29 allele was also absent from other populations and is present at a low frequency (AF = 0.0003) in the Indian population. HLA-B*15:27 was identified in Vietnam, Taiwan, Japan, with an AF >0.01 and Spain with AF <0.0001. HLA-B*27:07 was detected at a low frequency in India, Colombia, Spain, and South Africa. The HLA-B*46:01 allele was detected in Russia, Mexico, Vietnam, Taiwan, and Japan.

Sixty-six Brazilian individuals (17.1%) presented the HLA-DQB1*06:02 allele (AF=0.08938), 47 individuals (12.2%) carry the HLA-DRB1*15:01 allele (AF=0.06477) and 32 individuals (8.29%) have both the HLA-DRB1*15:01 and HLA-DQB1*06:02 alleles. The population of other continents, except Oceania, also have these two HLA alleles (HLA-DRB1*15:01 and HLA-DQB1*06:02) with an AF >0.01. Also, 15 Brazilian individuals (3.88%) carry HLA-B*15:03 (AF = 0.02073), which is predicted to have the greatest capacity to present SARS-CoV-2 peptides to immune cells^14^. This allele was not found in the Asian population of Japan, South Korea, and Vietnam (Supplementary Data 3).

### In silico analysis of viral peptide-HLA class I and II binding affinity

To verify the potential for an HLA allele type to produce an antiviral response to SARS-CoV-2, we performed HLA binding affinity analyses to the SARS-CoV-2 proteome. We tested 42 HLA-A proteins, 77 HLA-B, 38 HLA-C, 60 HLA-DP (DPA1/DPB1), 145 HLA-DQ (DQA1/DQB1), 46 HLA-DRB1, 4 DRB3, 2 DRB4, and 6 DRB5.

The SARS-CoV-2 proteome was presented by a diversity of HLA alleles from classes I and II (Supplementary Table 2). The HLA proteins are predicted to bind a small proportion of all possible SARS-CoV-2 derived peptides with high affinity (on average 0.5% for HLA class I and 2% for HLA class II). Also, we found a small proportion of weak binders (on average, 1.5% for HLA class I and 8.2% for class II). Most of the HLA proteins do not bind either Class I (on average >96%) or class II (on average >89%) molecules (Supplementary data 4). Supplementary Data 5 shows a list of HLA strongest binders (>300 peptides bound at high affinity) of SARS-CoV-2 peptides. These are found in loci HLA-A, -B, -C, and DQ.

## Discussion

Accessing the genomic sequences of the general population is relevant to identify the genetic variability involved in the molecular mechanisms of infection^20^. Also, it is known that the admixed Brazilian population is underrepresented in large public databases^21,22^, and previous studies revealed variants present exclusively in Brazilian indivuduals^19,24^.

We studied 27 human COVID-19-related genes and the HLA region in two public genomic databases of admixed Brazilians (BIPMed, www.bipmed.org^19^; ABraOM http://abraom.ib.usp.br/^24^), and additional samples from individuals born in three different towns of south-eastern Brazil. We reported the variants and HLA alleles found in these samples and compared them with worldwide populations. We also reported variants constituting the COVID-19 risk core haplotype on locus 3p21.31, described as being inherited from Neanderthals^12^.

Previous studies showed that the *ACE2, TMPRSS2, CTSL, FURIN*, and *IL6* genes, as well as the HLA region, may be involved in SARS-CoV-2 infection^1-5,10^ and immune response^6-8,14,23,25^. Furthermore, variants on loci 3p21.31 and 9q34.2 (encompassing *SLC6A20, LZTFL1, FYCO1, CXCR6, XCR1, CCR9*, and *ABO*) have been associated with Spanish and Italian patients with COVID-19^11^, and different variants were found to affect the predisposition to life-threatening illness in patients with COVID-19 from different ancestries^9^.

The analysis of genetic variability in candidate genes for specific populations can help to identify individuals at a higher risk of infection or severe disease by constructing risk haplotypes, which can also provide therapeutic targets for the development of more effective treatments and the control of COVID-19^2,10^. Thus, in addition to investigating genetic variability in the 27 candidate genes, we extended our analysis to include HLA alleles, which influence immunological response to many infectious agents (updated on September 28^th^ 2020; https://covid19.who.int/; https://coronavirus.jhu.edu/map.html). This is the first comprehensive study of genetic variability of COVID-19 genes in admixed individuals from Latin America, a hard-stricken population in the COVID-19 pandemic^26^, both in terms of the number of infected individuals and the severity of disease leading to increased death rates. Indeed, in the USA, remarkable disparities of SARS-CoV-2 infection by ethnicity have been shown, with Hispanic/Latino and African American individuals presenting higher SARS-CoV-2 infection rates and risk mortality than ‘non-Hispanic white’ Americans^27-29^. Therefore, by looking at population genomics data, one may gain insights into disease-related variants, which could be disproportionally represented in specific populations^18,30-32^. Furthermore, by evaluating individuals with unknown information on SARS-CoV-2 infection, one can achieve the random distribution of these variants, allowing better estimates of the distribution of population allele frequencies.

We identified small differences in AF in the 395 candidate variants identified among Brazilian samples, strengthening the hypothesis that different genetic backgrounds could influence SARS-CoV-2 infection and behaviour in human host cells^2,10^. Furthermore, this study and previous works^2,10^ identified individuals who carry unique deleterious variants, which may influence gene function and could potentially lead to different responses to SARS-CoV-2 infection on an individual scale. However, the rather similar distribution of AFs among Brazilians and their ancestry populations (NFE/EUR and AFR), as well as other admixed Americans (AMR), and the fact that the unique variants identified in the Brazilian population are rare or ultra-rare, indicates that the admixed Brazilian genetic background is not sufficient to influence SARS-CoV-2 infection on a population scale.

Zeberg and Pääbo^12^ have shown that the major genetic risk factor for severe COVID-19 was inherited from Neanderthals^12^. This finding is important on a regional scale, since 4% of admixed Americans analysed by Zeberg and Pääbo^12^ (including 1533 Brazilian controls from the BRACOVID dataset) presented the core haplotype derived from Neanderthals. Interestingly, Campinas, Ribeirão Preto, and the BIPMed dataset showed only one risk allele, while Barretos and Belo Horizonte did not present any risk allele of the Neanderthal’s core haplotype reported. Therefore, if further studies demonstrate that the Neanderthal-derived region confers a risk to COVID-19, this information should be carefully evaluated in additional admixed Brazilian samples from different geographic areas.

Currently, there is no consensus regarding a possible association of HLA alleles and susceptibility to COVID-19. Ellinghaus et al.^11^ did not find any evidence of an association between HLA and COVID-19. On the other hand, HLA-DRB1*15:01, HLA-DQB1*06:02, and HLA-B*27:07 alleles were associated with Italian patients affected by an extremely severe or severe form of COVID-19^23^, and an increased frequency of HLA-C*07:29 and HLA-B*15:27 was detected in Chinese patients with COVID-19 in comparison to the Chinese control population^25^. Interestingly, the HLA-C*07:29 allele is absent from the Brazilian admixed samples included in the present study and in all populations used in the comparisons, except for individuals from India, where this allele was found at a low frequency (0.0003). On the other hand, the HLA-B*15:27 allele was identified in individuals from three Asian countries (Vietnam, Taiwan and Japan) with AF >0.01, and at a low frequency in Spain (0.0001), but absent from Brazilian samples. The HLA-B*27:07 allele found in Italian individuals with a severe manifestation of COVID-19 was also identified in India, Colombia, Spain, and South Africa, but not in populations from Asia and Oceania (countries that are less affected by COVID-19) and from Brazil. In contrast, the HLA-DQB1*06:02 is present in all populations surveyed in this study, including Brazilian individuals (17.1%), with the exception of individuals from Australia. Also, the HLA-DRB1*15:01 allele is present in all populations investigated in this study, including Brazilian individuals (12.2%), but not in individuals from Australia and Peru. Interestingly, 8.29% of Brazilian individuals carry both the HLA-DRB1*15:01 and HLA-DQB1*06:02 alleles.

Furthermore, the HLA-A, -B, -C, and DQ loci show haplotypes that are strong binders of SARS-CoV-2 peptides in the Brazilian samples, especially for the HLA-A locus (20 alleles, Table 3). When comparing different populations, we found marked variability in the frequency of the different HLA alleles putatively associated with the severe manifestation of COVID-19, such as HLA-DRB1*15:01, HLA-DQB1*06:02, and HLA-B*27:07 alleles^23^. Overall, 10% of Brazilian individuals carried at least two of the alleles associated with the severe manifestation of COVID-19. Interestingly, the same alleles were absent from individuals from Australia. Variability in the frequency of HLA alleles previously associated with COVID-19 highlights the importance of considering ethnic and geographic origin when performing studies investigating the role of HLA alleles and disease. Thus, it seems likely that different population-specific haplotypes may be associated with an increased risk of severe disease in different populations.

**Table 3.** List of HLA strongest binders (>300 peptides bound at high affinity) of SARS-CoV-2 peptides and frequency in the Brazilian sample.

| HLA | HLA alleles | Allele frequency |
| --- | --- | --- |
| HLA -A | A*01:01 | 0.10233 |
|  | A*11:01 | 0.04145 |
|  | A*11:67 | 0.00130 |
|  | A*23:01 | 0.03756 |
|  | A*23:17 | 0.00389 |
|  | A*24:02 | 0.10104 |
|  | A*24:03 | 0.00259 |
|  | A*24:05 | 0.00130 |
|  | A*25:01 | 0.00389 |
|  | A*26:01 | 0.02979 |
|  | A*26:02 | 0.00130 |
|  | A*26:08 | 0.00130 |
|  | A*29:01 | 0.00259 |
|  | A*29:02 | 0.04534 |
|  | A*29:119 | 0.00130 |
|  | A*30:02 | 0.02591 |
|  | A*30:04 | 0.00259 |
|  | A*34:02 | 0.00777 |
|  | A*36:01 | 0.00389 |
|  | A*80:01 | 0.00130 |
| HLA-B | B*15:08 | 0.00130 |
|  | B*15:11 | 0.00130 |
| HLA-C | C*03:02 | 0.00389 |
|  | C*07:02 | 0.05699 |
|  | C*07:50 | 0.00130 |
|  | C*14:02 | 0.02979 |
|  | C*14:03 | 0.00259 |
| HLA-DQ | DQA1*0201-DQB1*0402 | 0.00259 |
|  | DQA1*0301-DQB1*0402 | 0.00130 |
|  | DQA1*0303-DQB1*0401 | 0.00130 |
|  | DQA1*0303-DQB1*0402 | 0.00130 |

In conclusion, we found rare variants in three COVID-19-related genes that are present only in the Brazilian dataset and are predicted to affect protein function. Furthermore, we identified HLA alleles previously associated with COVID-19 immunological response and 31 HLA alleles predicted as strong binders to SARS-CoV-2 peptides at loci -A, -B, -C, and DQ, which indicates the importance of further investigation on the role of HLA haplotypes as modulators of response to infection to SARS-CoV-2. Although the variants predicted to affect protein function in COVID-19-related genes are rare in admixed Brazilians (varying from 0.0001 to 0.0032), these also emerge as candidates for modulating response to infection by the SARS-CoV-2 in the Brazilian population. Furthermore, our study suggests the utility of population genomic studies in the context of precision health to stratify risk for infection disorders.

## Methods

### Subjects

We evaluated exomes from 257 individuals from the BIPMed dataset^19^, 609 from the ABraOM dataset^24^, and an additional 88 exomes from individuals born in three towns in south-eastern Brazil: Barretos (N=30), Ribeirão Preto (N=30), located in the state of São Paulo, and Belo Horizonte (N=28) the capital of the state of Minas Gerais. Among the BIPMed individuals, 193 had information about their city of birth available. The HLA region was sequenced in 386 individuals, including the 257 from BIPMed, the 88 additional exomes, and an additional 41 individuals (22 from southeast Brazil). We signed terms of data privacy to obtain permission to use the raw data from BIPMed and ABraOM public databases and use raw data of the 88 exomes from Barretos, Ribeirão Preto, and Belo Horizonte. This study was approved by the University of Campinas Research Ethics Committee (UNICAMP, Campinas, São Paulo, Brazil). All methods were performed according to the relevant guidelines and regulations.

### Exome analysis

Whole exome data were stored in variant call format (VCF) files built-in GRCh37/hg19 assembly. Gene regions were extracted by *vcftools*^33^ based on the position reported in Ensembl GRCh37 Release 101^34^ (Supplementary Table 1). Variant consequences were annotated from each gene region by ANNOVAR software (version 2019Oct24)^35^, using the following flags: *-otherinfo* (to include Brazil AF); *-onetranscript*; *-buildver hg19*; *-remove*; *-protocol refGene,gnomad211_exome*; *ALL*.*sites*.*2015_08*; *EUR*.*sites*.*2015_08*; *AFR*.*sites*.*2015_08*; *AMR*.*sites*.*2015_08*; *EAS*.*sites*.*2015_08*; *SAS*.*sites*.*2015_08*; *-operation g,f*; and *-nastring*. ANNOVAR software provides allele frequency (AF) information from African/African-American (AFR/AFA), Latino/admixed American (LAT/AMR), East Asian (EAS), non-Finish European (NFE), and South Asian (SAS) populations from gnomAD exome dataset^21^, as well as sub-Saharan Africans (AFR), Europeans (EUR), admixed Americans (AMR), east Asians (EAS), and south Asians (SAS) from 1KGP phase 3 dataset^22^. In addition, we annotated variants which were not identified by ANNOVAR using Variant Effect Prediction (VEP) algorithm^36^, with the following parameters: *--buffer_size 500*; *--canonical*; *--distance 5000*; *--species homo_sapiens*; *--symbol*.

To evaluate whether regional variability is observed among Brazilian samples, we separated the samples based on the city in which individuals were born, including 32 individuals from Campinas extracted from the BIPMed dataset.

### In silico *prediction analysis*

To predict the impact on protein function of the non-synonymous variants identified, we applied the following computer algorithms, which is currently recommended by the ACMG/AMP guidelines: PANTHER^37^, MutationTaster^38^, Condel^39^, PROVEAN^40^, PolyPhen2^41^, Sort Intolerant from tolerant (SIFT)^42^, Align Grantham Variation/ Grantham Difference score (GVGD)^43^, Combined Annotation Dependent Depletion (CADD)^44^, PhD-SNPg^45^, Functional Analysis through Hidden Markov Models (FATHMM)^46^, SNPs&GO^47^, and MutPred2 (http://mutpred.mutdb.org).

For Align-GVGD, we classified the variants based on the graded classifier with a cut-off of C35 or higher for deleterious classification. For CADD, we used the PHRED-like score with a cut-off of 20, below which the variants were classified as benign and otherwise deleterious. For MutPred2, we considered a score threshold of 0.50 for pathogenicity. For all other algorithms, we considered the classification provided as an output.

To access the impact of mutations on protein dynamics and stability, we used the DynaMut server (http://biosig.unimelb.edu.au/dynamut/)^48^. The server requires an input file of protein structure in PDB format or by providing the four-letter accession code for any entry on the Protein Data Bank database (PDB; http://wwpdb.org). The code for the *FURIN* gene used was 5jxg. The other proteins are not available in the PDB database to be tested.

### HLA analysis

We sequenced 11 HLA Loci (HLA-A, HLA-B, HLA-C, HLA-DRB1, HLA-DQB1, HLA-DPB1, HLA-DQA1, HLA -DPA1, HLA-DRB3, HLA-DRB4, HLA-DRB5) in 298 samples using NGSgo^®^ panels (GenDx, Utrecht, The Netherlands). The DNA libraries were loaded onto a MiSeq Sequencer (Illumina Inc., San Diego, CA, USA), and the data were analysed with the NGSengine v.2.16.2 software (GenDx, Utrecht, The Netherlands). We determined the HLA alleles from the remaining 88 exomes using the HLA-HD (HLA typing from High-quality Dictionary) tool v.1.3.0 ^49-51^. The IPD-IMGT/HLA database release 3.40.0 ^52^ was used as a reference. Even though we obtained results with six- and eight-digit precision, we restricted the results to four-digit accuracy to compare with published data. HLA allele frequencies were calculated by Arlequin v.3.5.2.2 software ^53^.

### In silico analysis of viral peptide-HLA class I and II binding affinity

We performed *in silico* analysis of viral peptide-HLA class I and II binding affinity across HLA proteins found in our population for the entire SARS-CoV-2 proteome. All HLA-A, -B, -C alleles were selected to assess the peptide-binding affinity of their corresponding proteins HLA-A, HLA-B, HLA-C, respectively. The HLA-DR is represented by HLA-DRA/DRB1 dimer. Since HLA-DRA is considered monomorphic, we just used the HLA-DRB1. The HLA-DP and DQ are represented by the HLA-DPA1/DPB1 dimer and HLA-DQA1/DQB1 dimer, respectively.

FASTA-formatted protein sequence data was retrieved from the National Center of Biotechnology Information (NCBI) database (https://www.ncbi.nlm.nih.gov/genome/browse#!/overview/Sars-cov-2). The follow eleven protein viral product was used in the analyses: ORF1ab (YP_009724389.1), Surface Glycoprotein (S) (YP_009724390.1), ORF3a (YP_009724391.1), Envelope (E) (YP_009724392.1), Membrane Glycoprotein (M) (YP_009724393.1), ORF6 (YP_009724394.1), ORF7a (YP_009724395.1), ORF7b (YP_009725318.1), ORF8 (YP_009724396.1), Nucleocapsid (N) (YP_009724397.2), and ORF10 (YP_009725255.1).

We k-merised these sequences into 8- to 12-mers to assess HLA class I-peptide binding affinity and into 15-mers to assess HLA class II binding affinity across the entire proteome. Predictions for HLA were performed using different HLA alleles found in our population with netMHCpan v4.1 for class I ^54^ and NetMHCIIpan - 3.2 for class II ^55^.

### HLA allele and haplotype frequencies of other populations

HLA frequency data were obtained from the Allele Frequency Net Database (http://www.allelefrequencies.net/) ^56^ for 10 distinct populations that are most and least affected by COVID-19. We checked the HLA of the populations that occupy the top 10 positions (USA, India, Brazil, Russia, Colombia, Peru, Spain, Mexico, Argentina, South Africa) and those that were less affected (Australia, Vietnam, Taiwan, Japan, and South Korea) (accessed on 04/24/2020, https://www.worldometers.info/coronavirus/) according to the availability of this data in the Allele Frequency Net Database.

## Data availability

BIPMed raw dataset that supports this study’s findings is available in EVA repository/PRJEB39251, https://www.ebi.ac.uk/eva/?eva-study=PRJEB39251. ABraOM raw dataset that supports the results of this study is available from ABraOM (http://abraom.ib.usp.br/).

## Acknowledgments

This work was supported by the Fundação de Amparo à Pesquisa do Estado de São Paulo (FAPESP, grant number 2013/07559-3). RS was supported by FAPESP (grant number 2019/08526-8). TKA is supported by FAPESP (grant number 2017/01900-6). IL-C is supported by CNPq (grant number 311923/2019-4).

## Competing interests

The authors declare no competing interests.

## Author contributions

RS contributed with the study design, conceptualization, data acquisition, analysis, and paper writing; TKA contributed with HLA sequencing, analysis, *in silico* prediction analysis, and writing of the paper; MCG contributed with *in silico* prediction analysis and paper writing; CSR contributed with public data acquisition and processing; MN and MZ contribute with public data acquisition; LD and MACB contributed with Belo Horizonte data acquisition and sample information; VLV contributed with Barretos data acquisition and sample information; WAS contributes with Ribeirão Preto data acquisition and sample information; ILC contributed with project conceptualization and served as principal investigators. All authors reviewed the manuscript.

## Notes

### Competing Interest Statement

The authors have declared no competing interest.

