## Supplementary tables for "Genetic variability in COVID-19-related genes in the Brazilian population"

**Supplementary Table 1.** Gene regions used to extract variants from VCF files, including the number of variants and coding variants per gene.

| Gene | Chr | Start (bp) | End (bp) | Number of variants | Number of coding variants* |
|---|---|---|---|---|---|
| *STAT1* | 2 | 191833762 | 191878976 | 411 | 6 |
| *SLC6A20* | 3 | 45796942 | 45838027 | 351 | 18 |
| *LZTFL1* | 3 | 45864808 | 45957534 | 447 | 7 |
| *CCR9*† | 3 | 45927996 | 45944667 | 131† | 5 |
| *FYCO1* | 3 | 45959396 | 46037316 | 681 | 67 |
| *CXCR6* | 3 | 45982425 | 45989845 | 44†† | 3 |
| *XCR1* | 3 | 46058516 | 46069234 | 88 | 2 |
| *TLR3* | 4 | 186990309 | 187006252 | 142 | 19 |
| *IL6* | 7 | 22765503 | 22771621 | 92 | 7 |
| *CTSL* | 9 | 90340434 | 90346308 | 99 | 11 |
| *ABO* | 9 | 136125788 | 136150617 | 493 | 31 |
| *IRF7* | 11 | 612555 | 615999 | 100 | 25 |
| *UNC93B1* | 11 | 67758575 | 67771593 | 201 | 21 |
| *STAT2* | 12 | 56735381 | 56754058 | 198 | 14 |
| *TBK1* | 12 | 64845840 | 64895899 | 355 | 16 |
| *IRF9* | 14 | 24630422 | 24635774 | 72 | 3 |
| *TRAF3* | 14 | 103243816 | 103377837 | 987 | 8 |
| *FURIN* | 15 | 91411822 | 91426688 | 236 | 23 |
| *TICAM1/TRIF* | 19 | 4815936 | 4831754 | 228 | 24 |
| *IRF3* | 19 | 50162826 | 50169132 | 98 | 22 |
| *IFNAR2* | 21 | 34602231 | 34636831 | 336 | 13 |
| *IFNAR1* | 21 | 34696748 | 34732129 | 273 | 12 |
| *TMPRSS2* | 21 | 42836478 | 42903043 | 765 | 21 |
| *ACE2* | X | 15579156 | 15620271 | 221 | 12 |
| *NEMO/IKBKG* | X | 153769419 | 153796804 | 123 | 5 |
| **Total** | | | | **7172** | **395** |

*missense variants, frameshifts, stop gained, and splicing sites; †overlap with *LZTFL1*; ††overlap with *FYCO1*; Chr= chromosome.

**Supplementary Table 2.** Number of peptides binders for HLA alleles.

| Proteins | Class I | Class II |
|---|---|---|
| ORF1ab | 35435 | 7082 |
| Orf8 | 560 | 107 |
| E | 330 | 61 |
| M | 1065 | 208 |
| N | 2050 | 405 |
| orf3a | 1330 | 261 |
| orf6 | 260 | 47 |
| orf7a | 560 | 107 |
| orf7b | 170 | 29 |
| orf10 | 145 | 24 |
| S | 6320 | 1259 |
| **Total** | **48225** | **9590** |
